# RUVBL1 and RUVBL2 are druggable MYCN regulators in neuroblastoma

**DOI:** 10.1101/2024.10.03.616410

**Authors:** Joachim Tetteh Siaw, Arne Claeys, Wei-Yun Lai, Marcus Borenäs, Elien Hilgert, Sarah-Lee Bekaert, Ellen Sanders, Irem Kaya, Jo Van Dorpe, Frank Speleman, Kaat Durinck, Bengt Hallberg, Ruth H. Palmer, Jimmy Van den Eynden

## Abstract

High-risk neuroblastoma is characterized by *MYCN* amplification and high *MYCN* or *MYC* gene expression. These patients have a poor prognosis and there is an urgent need for more effective drugs. While strategies to develop inhibitors that directly target the MYC proteins have remained largely unsuccessful, recent preclinical studies have identified ATR, a key protein of the DNA damage response, as a promising alternative therapeutic target. Here we identified a strong RUVBL1 and RUVBL2 signature in transcriptomics data derived from different *MYCN*-driven mice tumors treated with ATR inhibitors. The RUVBL proteins form a complex with ATPase activity that has broad cellular functions and we demonstrate that pharmacological inhibition of this protein complex results in a strong reduction of MYC signaling, cell cycle arrest, DNA damage and apoptosis. We confirmed the association with *MYCN* and identified the *RUVBL* genes as independent prognosticators in human primary neuroblastoma data.

## INTRODUCTION

Neuroblastoma (NB) is the most common cancer in infancy. It arises from neural crest derived immature sympathoblasts and occurs along the sympathetic chain ganglia and in the adrenal gland^1,2^. Despite the improvement in survival over the years, more than 50% of high-risk patients relapse despite intensive multimodal therapies^3–5^. Additionally, severe life-threatening toxicities occur in many high-risk patients^5,6^. Hence, safer and more effective drugs are urgently needed.

NB cells are genomically characterized by recurrent patterns of distinct segmental chromosomal aberrations including deletions in 1p and 11q, gains in 2p and 17p and *MYCN* amplification^7–11^. *MYCN* amplification and high *MYC* expression are associated with the majority of high-risk NB cases^12–15^. While the inhibition of the MYCN (also known as n-Myc) and MYC (known as c-Myc) proteins would be a highly effective therapeutic strategy, the development of such inhibitors has proven challenging^16–18^.

Recent research identified ATR, CHK1 and other proteins involved in the NB cell’s DNA damage response (DDR) as promising therapeutic targets in high-risk NB^19–22^. Here, we identified RUVBL1 (also known as Pontin) and RUVBL2 (known as Reptin) as key mediators in this therapeutic response. RUVBL1 and RUVBL2 (commonly referred to further as RUVBL1/2) are members of the AAA+ (ATPase associated with diverse cellular activities)-family of ATPases and form heterohexameric or heterododecameric protein complexes that are involved in a broad range of cellular activities including transcriptional co-activation, epigenetic regulation of gene expression, cell proliferation, DNA repair, regulation of telomerase activity and senescence^23–29^. We experimentally demonstrate that pharmacological inhibition of RUVBL1/2 results in reduced MYC and MYCN activity, DNA damage and apoptosis in several NB cell lines and clinically validated these findings in large NB patient data sets by showing that *RUVBL1/2* are strong prognosticators, independent of *bonafide* biomarkers.

## RESULTS

### Identification of the RUVBL1/2 complex as a putative therapeutic target in neuroblastoma

We previously demonstrated that therapeutic inhibition of ATR in genetically engineered ALK/MYCN-driven NB mice models is a highly effective treatment strategy^19,21^. To identify other putative treatment targets for NB, we mined our published ATR inhibitor-treated mice tumor RNA-Seq data for key transcriptional regulators involved in the observed treatment effects. A gene set enrichment analysis (GSEA) was performed using published transcriptional regulator targets that were identified using Chip-Seq on mouse embryonic stem cells. The targets of the transcriptional co-factor RUVBL2 was the most significantly depleted gene set in *Th-MYCN-ALK^F^*^178^*^F^*mice treated for 3 days with the ATR inhibitors elimusertib (25 mg/kg; NES = -2.1, *P* = 9.66e-57) and ceralasertib (25 mg/kg; NES = -2.7, *P = 1.56e-97*), as well as in elimusertib-treated *Th-MYCN-ALKAL2* mice (NES *=* -2.1, *P = 1.17e-43*; Fig. 1A-B). RUVBL2 forms a protein complex with RUVBL1 and, as expected, strong depletions were also observed when focusing on the targets of RUVBL1 (NES = -1.9 or lower, *P* = 1.5e-13 or lower, Fig. 1A-B). Apart from the downregulation of the RUVBL targets, we also observed a significant downregulation of the *RUVBL* genes themselves in all 3 experimental conditions (P < 0.001; Fig. 1C). These results suggest that the treatment effects observed upon ATR inhibition could be at least partially mediated through RUVBL downregulation, identifying the latter as a putative (combinatorial) drug target for neuroblastoma.

**Figure 1.**
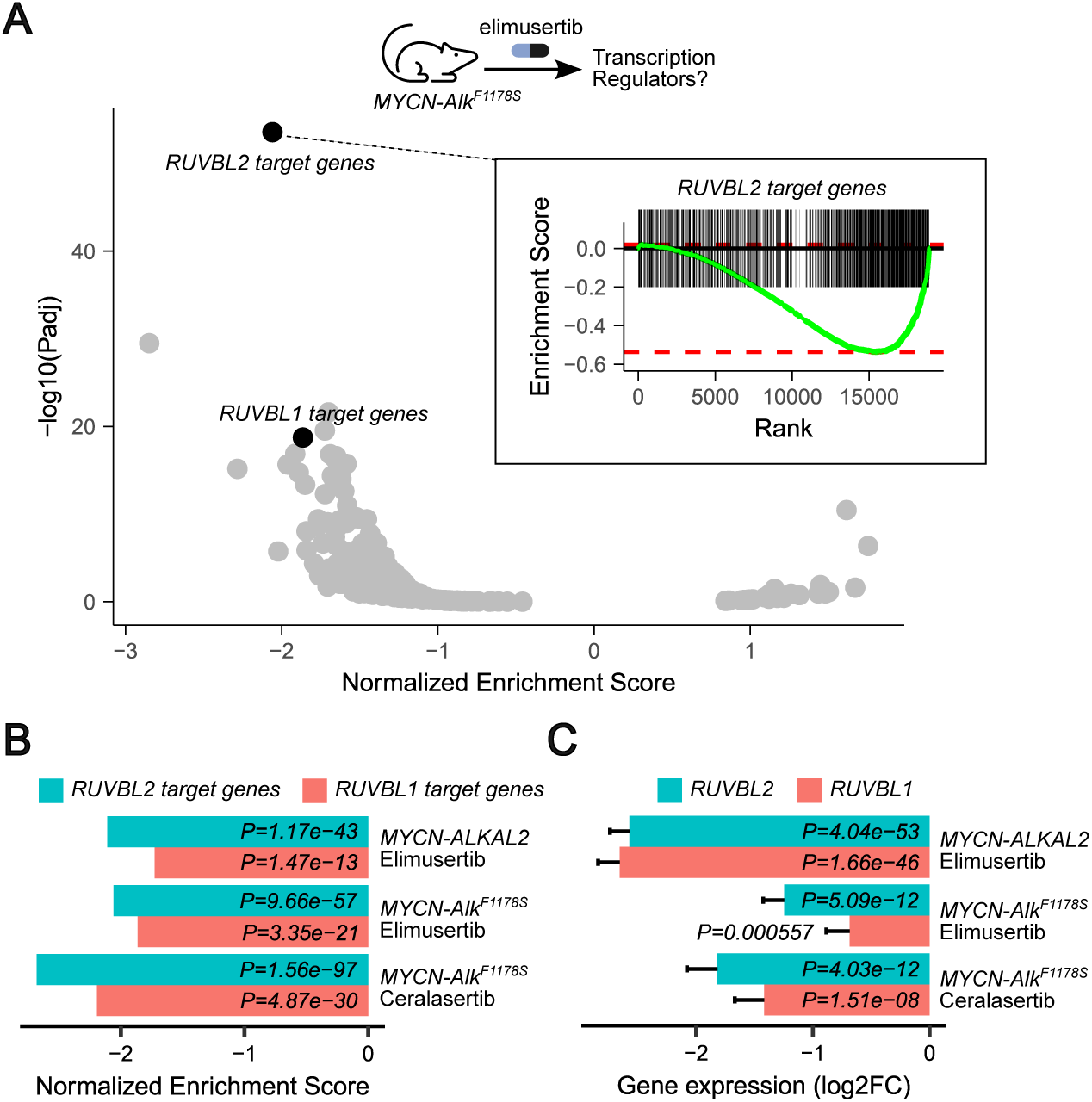
Transcriptional regulator gene set enrichment analysis on NB mice treated with ATR inhibitors. Differential gene expression (DGE) data obtained from *MYCN-ALK^F^*^1178^*^S^* and *MYCN-ALKAL2* driven NB mice tumors treated with the ATR inhibitors elimusertib and ceralasertib (both 25 mg/kg per day and after 3 days of treatment) were obtained from previous studies^19,21^. **A.** Volcano plot showing GSEA results performed using transcriptional regulator target genes on elimusertib treated *MYCN-ALK^F^*^1178^*^S^* tumors as indicated. Running score plot for RUVBL2 target genes shown on inset. **B.** Bar plot showing normalized enrichment scores (NES) and corresponding *P* values for RUVBL1 and RUVB2 target genes in 3 experimental conditions as indicated. **C.** Bar plot showing differential *RUVBL1* and *RUVBL2* gene expression results (log2 fold change and *P* values) in 3 experimental conditions as indicated.

### RUVBL1 and RUVBL2 are druggable dependency genes in neuroblastoma cell lines

To find support for functional roles of *RUVBL1/2* in NB, we first analyzed publicly available gene dependency scores of 34 NB cell lines, with negative scores indicating gene dependency^30^. Interestingly, both *RUVBL1* (median score = -1.83) *and RUVBL2* (median score = -1.75) had significantly lower scores than a previously identified set of 1910 essential genes (median score = -1.00; *P* < 0.0001; unpaired Wilcox test; Fig. 2A).

**Figure 2.**
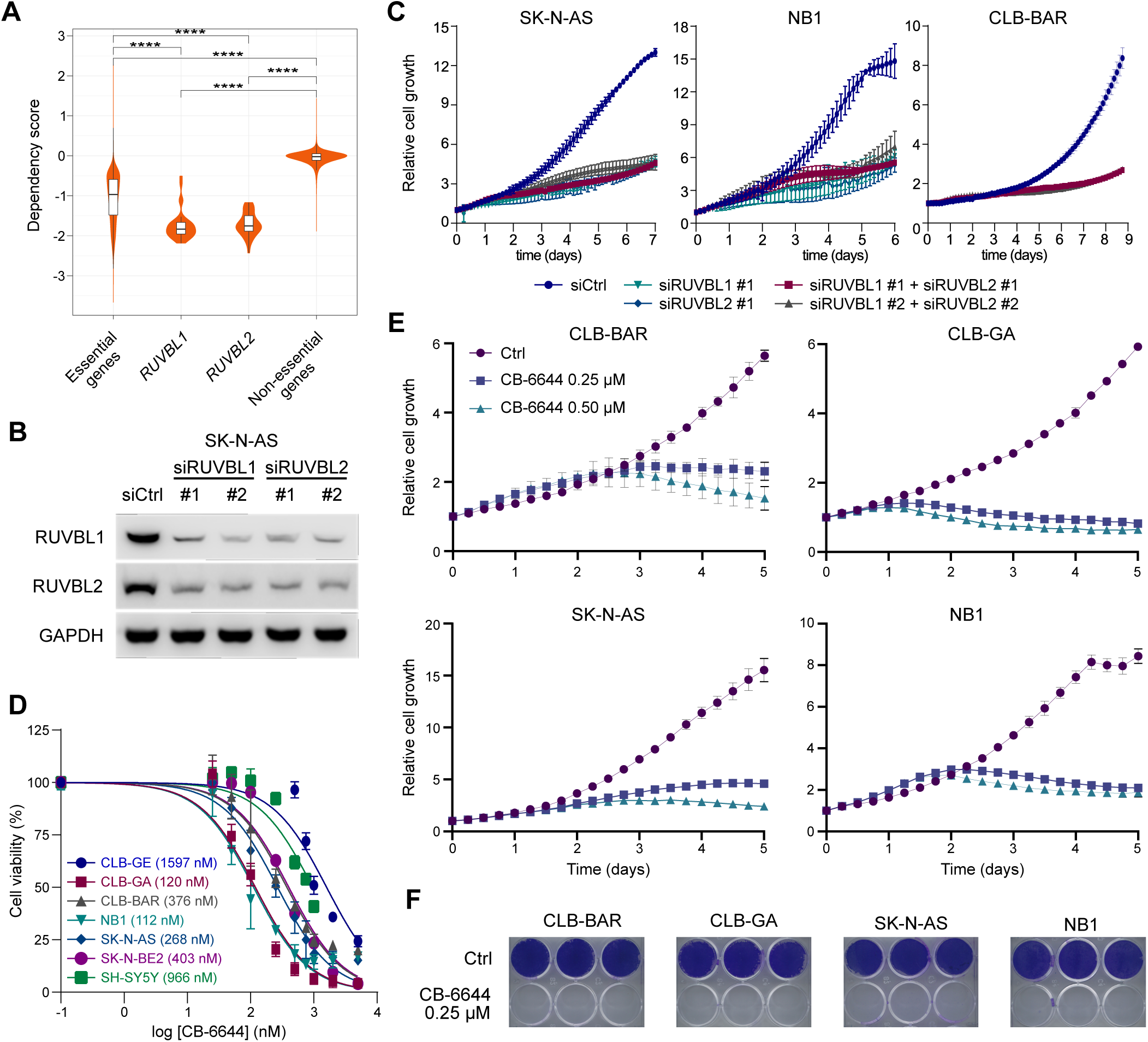
Pharmacological inhibition of RUVBL1/2 ATPase activity with CB-6644, in NB cell lines. **A.** Violin plots comparing DepMap *RUVBL1* and *RUVBL2* dependency scores from 34 NB cell lines to sets of essential and non-essential genes. **** *P* value < 0.0001, two-sided Wilcoxon rank sum test. **B.** Western blot showing transient siRNA-mediated knockdown of *RUVBL1* and *RUVBL2* after 3 days of transfection. Scrambled siRNA was used as control (siCtrl). **C, E.** Time-dependent effect of siRNA-mediated knockdown of *RUVBL1* and/or *RUVBL2* (**C**) and CB-6644 treatment (250 nM or 500 nM, as indicated; **E**) on NB cell line proliferation. Cell proliferation was monitored live by scanning for cell confluency at regular time intervals with IncuCyte® S3 system. Cell growth was normalized relative to the first scan at time zero. Results are mean +/-SEM of 3 independent biological replicates. **D.** Cell viability dose dependency curves after treatment with the RUVBL1/2 inhibitor CB-6644 in 7 different NB cell lines, as indicated. Mean IC50 values are indicated for each cell line (2 to 6 biological replicates). Cell viability was determined by resazurin assay. **F.** Long-term (14 days) effect of CB-6644 (250 nM) on NB cell growth. Cells were stained with crystal violet.

To further evaluate the specific cellular effects of RUVBL1/2 downregulation, we carried out transient knockdown using two independent pairs of small interfering RNAs (siRNAs) in the CLB-BAR, NB1 and SK-N-AS NB cell lines. Each siRNA efficiently knocked down its target, RUVBL1 or RUVBL2 and, interestingly, knockdown of RUVBL1 also resulted in downregulation of RUVBL2 and vice versa (Fig. 2B). We monitored cell growth for 1 week and observed a decreased cell number after 3-5 days of treatment as compared to control conditions in all studied cell lines (Fig. 2C).

Based on these results, we aimed to determine whether RUVBL is a putative therapeutic target in NB and treated seven different NB cell lines, with diverse genetic backgrounds, with a recently reported small molecule inhibitor, CB-6644, which specifically blocks the ATPase activity of the RUVBL1/2 complex^31^. We first determined the sensitivity of the different cells to CB-6644 using a resazurin cell viability assay (Fig. 2D) and then assessed cell growth upon CB-6644 (0.25 µM and 0.50 µM) treatment for the 4 most sensitive cell lines (CLB-BAR, CLB-GA, SK-N-AS and NB1). This treatment resulted in a complete block of NB cell growth starting 1-3 days after treatment initiation, depending on the cell line (Fig. 2E-F).

### The transcriptomic response upon pharmacological RUVBL inhibition is characterized by reduced MYC signaling in NB cells

To better understand the mechanisms underlying this pharmacological RUVBL inhibition, we performed RNA-Seq of the MYCN-driven CLB-BAR and MYC-driven SK-N-AS NB cells after treatment with CB-6644 (0.25 µM) for 24, 48 and 72 hours. The transcriptional response increased progressively over time and was more pronounced in CLB-BAR (1781 differentially expressed (DE) genes after 72h) as compared to SK-N-AS cells (989 DE genes; Suppl. Fig. 1A-B).

In CLB-BAR cells 627 genes were downregulated after 72h, including *MYCN*, *RRM2*, *RPS6*, *MCM5-7* and *NPM1* (Fig. 3A). A Hallmark GSEA indicated a strong enrichment for MYC target genes (*P_adj_* = 4.7e-50; Fig. 3B), MTORC1 signaling and the cell cycle-related targets of E2F transcription factors and G2/M checkpoints, strikingly similar to what we observed previously upon ATR inhibition with elimusertib (Fig. 3C)^19,21^. In SK-N-AS cells (517 genes downregulated after 72h), similar though overall weaker enrichments were observed as compared to CLB-BAR cells upon CB-6644 drugging (Fig. 3C). A Reactome pathway GSEA confirmed the enrichment for genes involved in cell cycle control, mainly in CLB-BAR cells. Additionally strong enrichments were observed for several RNA-related processes (e.g. RNA metabolism, translation, rRNA processing, nonsense-mediated decay) in both cell lines (Suppl. Fig. 1C; Suppl. Table 1).

**Figure 3.**
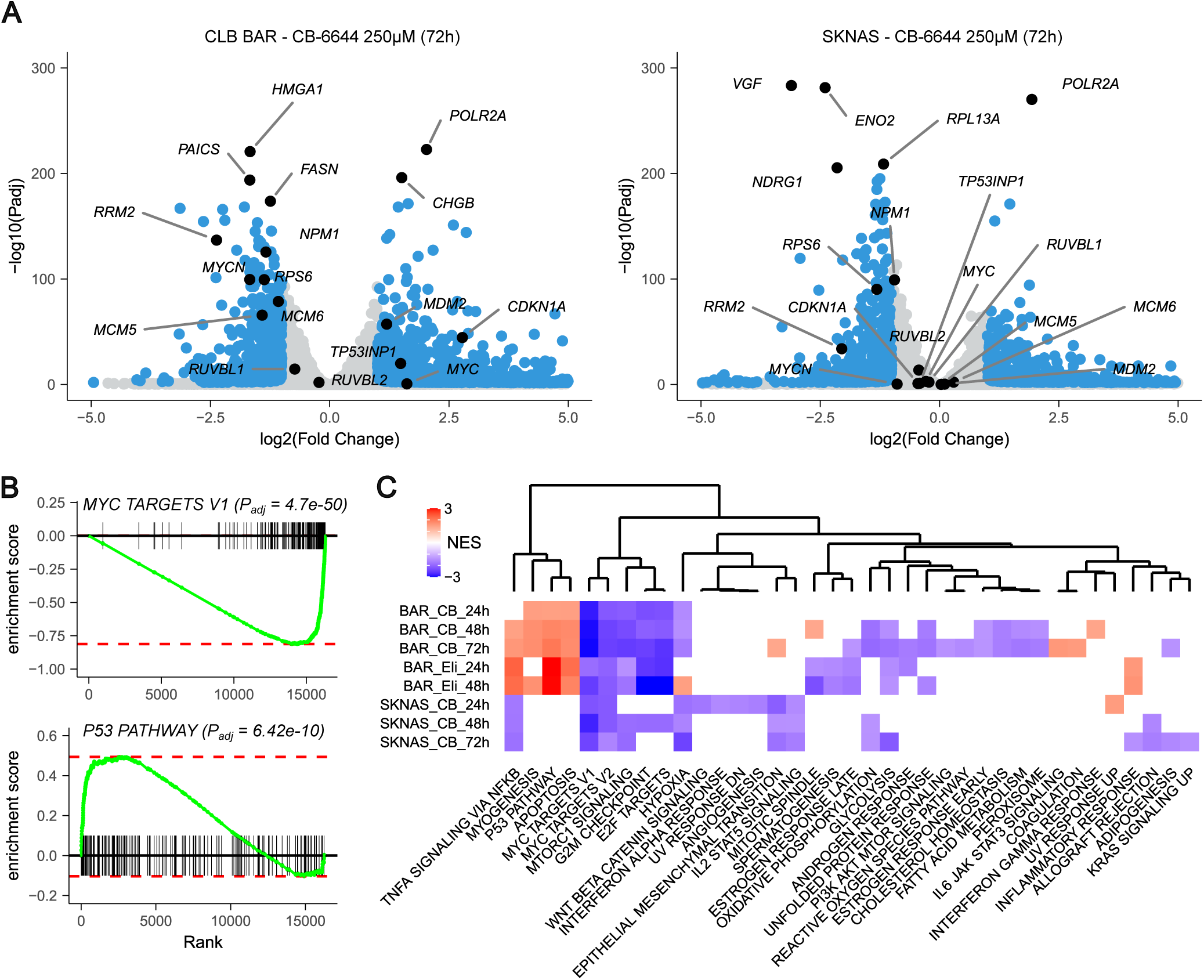
Transcriptomic response to CB-6644 treatment of NB cells. CLB-BAR and SK-N-AS NB cells were treated for 72h with CB-6644 250 nM and differential gene expression (DGE) was determined. **A.** Volcano plot for both cell lines as indicated. Differentially expressed genes (threshold log2FoldChange of ±1 at 1% FDR) indicated in blue with top up/downregulated and genes discussed in main text labelled. **B.** GSEA running score plots for 2 Hallmark gene sets in CLB-BAR cells as indicated. **C.** GSEA results with heatmap comparing normalized enrichment values (NES, color key) for both cell lines after treatment with CB-6644 (CB; 250 nM) and elimusertib (Eli; 50 nM) as indicated. Gene sets that were not significant (Padj < 0.05) were left blank and are not shown not significant for all analyzed conditions. Columns (gene sets) were hierarchically clustered as indicated in dendrogram. See Suppl. Table 1 for detailed DGE and GSEA results.

A different pattern was observed for the upregulated genes (1154 and 472 genes in CLB-BAR and SK-N-AS cells, respectively). While the Hallmark P53 (*P_adj_* = 6.4e-10) and apoptosis pathways were strongly enriched in CLB-BAR cells, similar to what we observed upon elimusertib treatment, these enrichments were remarkably absent in SK-N-AS cells (*P_adj_* > 0.05; Fig. 3C). Notably, *POLR2A* was the most upregulated and fastest responding gene in both cell lines (Fig. 3A, Suppl. Fig. 1B).

### Bidirectional regulation between MYCN and RUVBL1/2 in NB cells

We next aimed to validate this NB cellular response upon CB-6644 treatment. We first confirmed the significant downregulation of *MYCN* (CLB-BAR cells) and *MYC* (SK-N-AS cells) after 48h and 72h at the transcriptional level using quantitative PCR (Suppl. Fig. 2A). Similar to what we observed in the RNA-Seq data, the mRNA reduction was weaker in SK-N-AS (28% lower after 72h) as compared to CLB-BAR cells (61% lower). Western blotting confirmed this lower expression at the protein level in 4 different NB cell lines (Fig. 4A). Similar results were obtained after siRNA mediated knockdown of *RUVBL1* or *RUVBL2*, confirming *RUVBL1/2* specificity (Suppl. Fig. 3).

**Figure 4.**
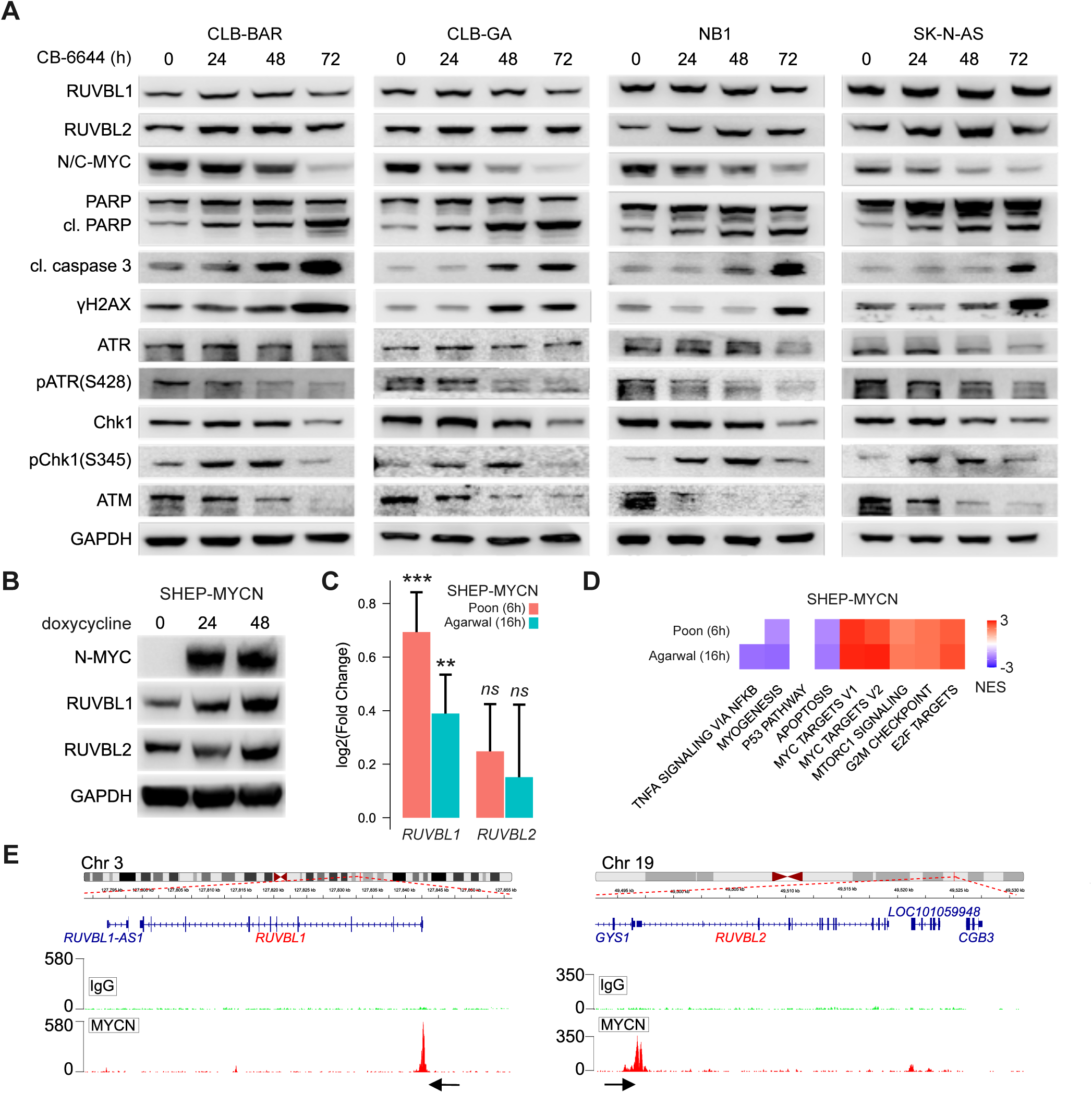
Experimental validation of transcriptomic responses upon CB-6644 treatment of NB cells. **A.** Western blots showing the effect of different CB-6644 (250 nM) treatment duration on different (phospho-)proteins in 4 NB cell lines as indicated. N/C-MYC denotes MYCN for CLB-BA, CLB-GA and NB1, and MYC for SK-N-AS and were detected by two independent antibodies. Cl, cleaved. **B.** Western blot showing the effect of doxycycline-induced *MYCN* overexpression on RUVBL1 and RUVBL2 protein expression in SHEP cells. **C-D.** Differential gene expression results of *MYCN* induction in SHEP cells of 2 independent RNA-Seq studies as indicated (time after induction between brackets). **C.** Bar plot showing *RUVBL1* and *RUVBL2* upregulation (log2 (Fold Change) +/-SEM). **, P < 0.01; ***, P < 0.001, ns, non-significant. **D**. Heatmap showing GSEA NES scores for the 2 main clusters as in Fig. 3C. **E.** CUT&RUN genome tracks showing MYCN binding peaks (red) at *RUVBL1* and *RUVBL2* promoter regions as indicated. IgG background signal shown in Green. Black arrows indicate direction of *RUVBL1* or *RUVBL2* transcription.

To determine whether this regulation could be bidirectional (i.e., MYCN regulating *RUVBL1/2* expression), we performed an RUVBL1/2 western blot before and after inducing *MYCN* in SHEP MYCN cells. Expression of RUVBL1, and to a lesser extent RUVBL2, was higher 24-48 hours after MYCN induction (Fig. 4B). We then queried the Cell Line Exploration web Application of NB (CLEAN^32^) and retrieved RNA-Seq data from 2 other, previously published studies on SHEP-MYCN cells, one after 6h of MYCN induction (Poon *et al*.^33^), the other after 16h (Agarwal *et al*.^34^). Both studies confirmed increased *RUVBL1* and *RUVBL2* expression after *MYCN* induction, although this was only significant for *RUVBL1* (P < 0.01 for both studies) and not for *RUVBL2* (Fig. 4C). Interestingly, a GSEA on these data identified an upregulation of the same cluster of gene sets (MYC targets, MTORC1 signaling, G2M checkpoints, E2F targets) that we observed to be downregulated upon pharmacological RUVBL inhibition (Fig. 4D). Additionally, when analyzing previously published gene expression data of samples taken during different stages of mice NB tumor development, we noticed a developmental increase in *Ruvbl1* and *Ruvbl2* expression in homozygous *Th-MYCN* mice but not in wild-type mice (Suppl. Fig. 2B), supporting a role of RUVBL1/2 in mediating MYCN effects. To determine whether RUVBL1/2 are direct transcriptional targets of MYCN in NB, we performed a CUT and RUN experiment and confirmed MYCN binding at the promotor regions of both *RUVBL1* and *RUVBL2* (Fig. 4E). Taken together, our results suggest a bidirectional regulation between MYC and RUVBL1/2 in NB cells.

### Pharmacological RUVBL inhibition results in DNA damage, S-phase arrest and apoptosis in NB cells

Next to the downregulated MYC targets, the CB-6644-induced transcriptomic response was further characterized by increased apoptosis signaling and cell cycle alterations (i.e., downregulation of E2F targets and G2M checkpoints), mainly in CLB-BAR cells. This apoptosis induction was confirmed in all examined cell lines by elevated expression of cleaved PARP as well as cleaved caspase 3 (Fig. 4A). We further confirmed this apoptosis induction by increased caspase 3/7 activity as assessed by the Caspase-Glo 3/7 apoptosis assay in CLB-GA and SK-N-AS cell lines (Fig. S4A). Relatedly, when assessing the NB cell cycle using propidium iodide staining and flow cytometry upon treatment of CLB-GA and SK-N-AS cells with CB-6644, a significant prolongation of S-phase was observed in both CLB-GA (*P* = 0.013) and SK-N-AS cells (*P* = 0.0072) after 48h of treatment (Fig. S4B).

Given the putative role of RUVBL1/2 in mediating at least part of the therapeutic effects we previously observed upon inhibition of ATR (Fig. 1), in combination with the previously suggested PIKK interactions^35,36^, we examined whether RUVBL inhibition resulted in an alteration of the DNA damage response. After 48h-72h of CB-6644 treatment we observed decreased phosphorylation levels of pATR^S428^ as well as the main ATR target pChk1^S345^. This reduced ATR-CHK1 signaling coincided with increased expression of the DNA damage marker γH2AX as well as the transcriptomic, proteomic and functional indicators of apoptosis. Notably, this reduced DDR signaling was preceded by increased phosphorylation of pChk1^S345^ 24h after treatment and followed by reduced protein expression of ATM and to a lesser extent also ATR and CHK1 after 72h (Fig. 4A).

### *RUVBL1* and *RUVBL2* are *MYCN*-independent prognosticators in human neuroblastoma

Our previous results suggest that *RUVBL1* and *RUVBL2* are therapeutically targetable dependency genes in NB. To find clinical evidence for such a role we focused on different sources of primary NB tumors and data.

We first confirmed protein expression of RUVBL1 and RUVBL2 in primary NB cancer cells using immunohistochemistry (Fig. 5A). We then focused on a large set (n = 364) of previously published primary NB RNA-Seq and associated clinical data. Similar to what we observed in cell lines, *MYCN* and *RUVBL1/2* expression were positively correlated, both in *MYCN* amplified and in *MYCN* wild-type tumors (Pearson’s *r* between 0.18 and 0.44; Fig. 5B). Relatedly, *MYCN* amplified tumors had significantly higher *RUVBL1* (*P* = 9.6e-28) and *RUVBL2* (*P* = 2.0e-22) expression as compared to *MYCN* wild-type tumors (Fig. 5C). Additionally, significantly worse survival was observed between tumors with high (above median) expression of *RUVBL1* (*P* = 3.2e-22; 5y survival 51%) or *RUVBL2* (*P* = 7.4e-20; 5y survival 54%) as compared to tumors with low expression of these genes (5y survival 93% or higher; Fig. 5D). Remarkably, despite the strong correlation between *RUVBL1/2* expression and *MYCN* amplification state, high *RUVBL1* and *RUVBL2* expression were identified as strong and *MYCN* amplification-independent prognosticators of NB outcome. Indeed, a Cox multivariate proportional hazards regression that considered higher-than-median *RUVBL1* or *RUVBL2* expression together with *bonafide* prognostic NB biomarkers as covariates (i.e., tumor stage, age at diagnosis and *MYCN* amplification) indicated significantly elevated hazard ratios for both high *RUVBL1* (HR = 3.4*; P* = 3.0e-04) and high *RUVBL2 (*HR = 3.0*; P* = 4.3e-04) expression (Fig. 5E).

**Figure 5.**
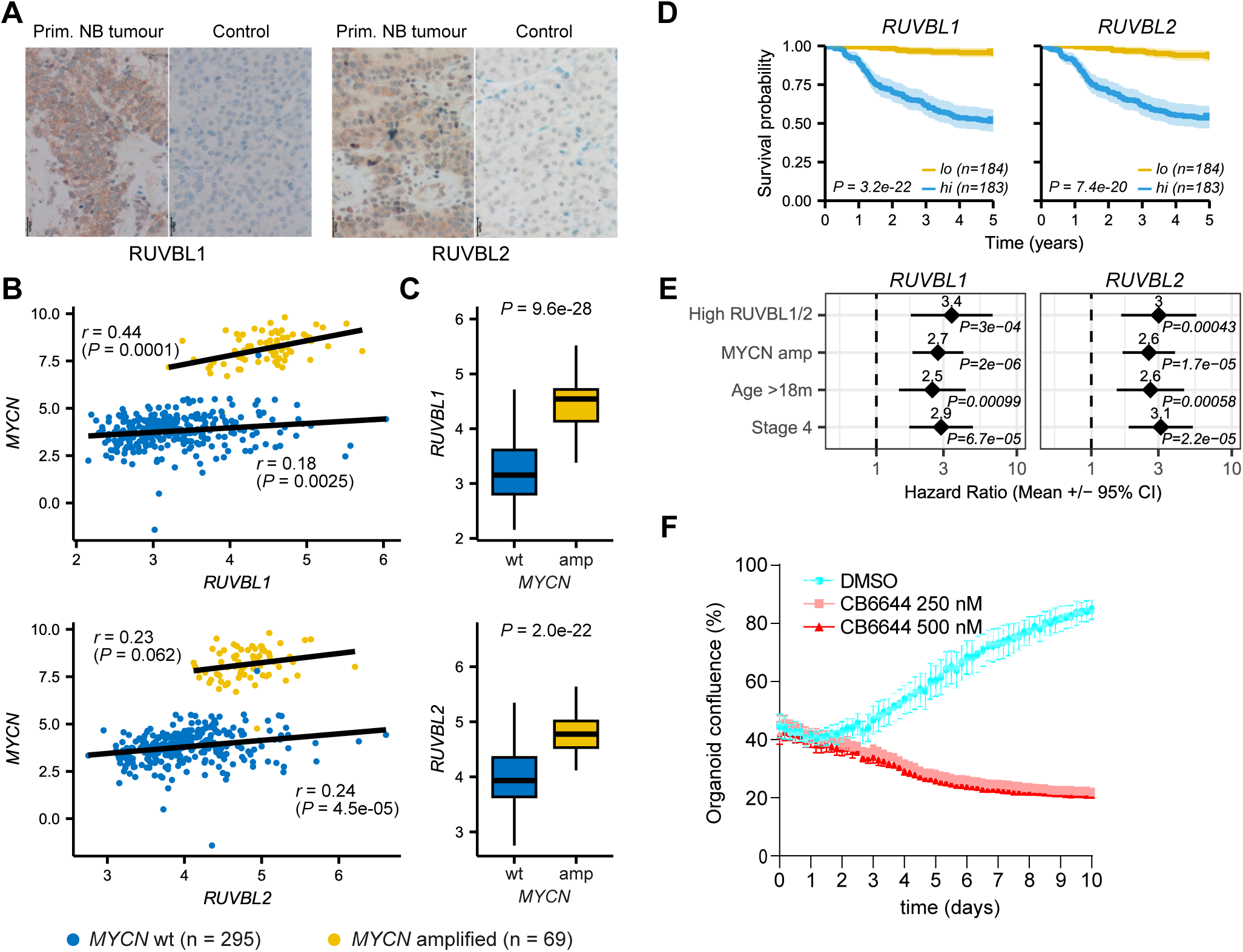
Clinical validation of the prognostic and putative therapeutic relevance of the *RUVBL* genes. **A.** Immunohistochemical staining for RUVBL1 and RUVBL2 in human NB tumor tissue sections as indicated. Normal pancreatic tissue was used as negative control for antibody specificity. Images are representative of 3 independent NB tumors and 2 independent pancreatic stained tissues. **B-E.** Analysis of publicly available primary neuroblastoma data (n=364). **B.** Correlation plots between *RUVBL1* (top), *RUVBL2* (bottom) and *MYCN* gene expression (log2 normalized counts) for *MYCN* amplified and *MYCN* wild-type tumors. Linear regression line and Pearson’s correlation coefficient indicated. **C.** Boxplots comparing *RUVBL1* (top) and *RUVBL2* (bottom) expression between *MYCN* amplified and *MYCN* wild-type tumors. P value calculated using two-sided unpaired Wilcoxon’s test. **D.** Kaplan-Meier survival plots comparing overall survival between patients with high and low *RUVBL1* and *RUVBL2* as indicated. High/low *RUVBL1/2* expression defined based on median gene expression. *P* value calculated using log rank test. **E.** Forest plots comparing hazard ratios +/-95% confidence intervals for 4 variables as indicated. Results were obtained using a Cox proportional hazards multivariate regression analysis. **F.** Time-dependent effect of CB-6644 (250 nM or 500 nM as indicated) on human NB PDX organoid growth. Organoid growth was monitored by scanning confluency at regular intervals with IncuCyte® Live Cell Analysis system. Results are mean +/- SD of three technical replicates. Graph is representative of 3 biological repeats with different organoid seeding densities.

Finally, we explored the therapeutic potential of CB-6644 RUVBL inhibition by treating organoids derived from a human high-risk, *MYCN* amplified NB patient. Similar to our cell line results, organoid growth was completely abrogated after treatment with CB-6644 250 nM or 500 nM (Fig. 5F). Altogether, these reinforce *RUVBL1/2* as dependency genes in NB pathogenesis.

## DISCUSSION

The DDR is a target of major interest in high-risk NB with several recent studies reporting highly promising results when therapeutically targeting key molecules such as ATR, AURKA or CHK1 in NB animal models^19–22^. In this study we discovered a strong *RUVBL1* and *RUVBL2* signature in transcriptomic data obtained from tumors treated with ATR inhibitors in *ALK*/*MYCN*-driven NB mice models. *RUVBL1* and *RUVBL2* form a protein complex with ATPase activity that can be inhibited by the small molecule inhibitor CB-6644. Based on a series of RNA-Seq and experimental validation experiments we demonstrated that CB-6644 inhibition of RUVBL1/2 activity resulted in reduced MYC signaling, apoptosis, cell cycle arrest and increased DNA damage. In primary NB data we observed a strong association between high *RUVBL1/2* expression and bad outcome, independent of the main risk factors (*MYCN* amplification status, age and stage). Our results suggest that RUVBL1/2 is a therapeutically targetable dependency gene and important prognosticator in NB.

One of the most striking signals we observed upon pharmacological RUVBL1/2 inhibition was the downregulation of MYC targets. These targets are regulated by the MYC family of transcription factors, which include MYCN (n-MYC) and MYC (c-MYC) and are amongst the most potent oncogenes in high-risk NB. Their lack of apparent surfaces for small molecule binding makes them difficult to target therapeutically^37^, and our results identify the RUVBL1/2 complex as an indirect therapeutic MYC target in NB, similar to recent findings in pancreatic cancer^38^. A positive correlation was found between *MYCN* and *RUVBL1/2* mRNA levels in NB cell lines as well as primary NB tumors, suggesting an interdependency at the transcriptional level, in line with previous reports^23,39^. Interestingly, we confirmed that this transcriptional regulation was bidirectional and demonstrated MYCN binding at *RUVBL1/2* promotor regions. Additionally, a recent study has identified RUVBL1/2 as a cofactor of MYC that is essential for chromatin association^38^, providing alternative explanations for the reduced MYC signaling upon RUVBL inhibition.

The transcriptomic response upon RUVBL1/2 inhibition was similar to what we observed upon ATR inhibition with a reduction of MYC, E2F, MTORC1 and G2M checkpoint signaling and increased (p53-related) apoptosis signaling. The RUVBL1/2 complex is a critical component of the HSP90-interacting chaperone-like complex R2TP and this complex is involved in the stabilization of PIKKs^40^. This stabilization function is in line with the reduced MTORC1 signaling and decreased ATR and ATM protein expression we observed after 24-48h of RUVBL inhibition. One of the major effectors of MTORC1 signaling is the ribosomal protein S6 kinase (S6K). This protein controls protein synthesis through phosphorylation of the ribosomal protein S6 (*RPS6* gene) which largely explains the reduced rRNA processing, RNA metabolism, and translation activities we observed in our transcriptome data upon RUVBL inhibition. Interestingly, our study indicated a strong and consistent downregulation of *RPS6* and in a previous protein kinase analysis we found strongly reduced kinase activity of several related ribosomal kinases upon ATR inhibition^21,32^.

The transcriptional response could be (partially) mediated via the RNA polymerase II (Pol II) complex. Indeed, the RUVBL1/2 complex has been shown to be essential for both the assembly and the regulation of the RNA polymerase II (Pol II) complex, explaining a broad range of transcriptional activities^23,40^. In this regard, it’s striking that the Pol II encoding *POLR2A* gene was the earliest and strongest upregulated gene, putatively representing a compensatory effect of decreased Pol II activity. Replication stress is a hallmark of high-risk NB and, interestingly, recent evidence suggests that RUVBL1 is involved in MYC multimerization near stalled replication forks, protecting them against Pol II-mediated replication stress and reducing double-strand break formation^41^.

A role of RUVBL has been suggested in many cancer types, including colorectal cancer, breast cancer, lung cancer and many others^35,38,40,42,43^. In general, these studies found tumor-promoting effects of *RUVBL1* and/or *RUVBL2* and a link between high RUVBL1/2 expression and poor clinical outcome. Our study demonstrates that that RUVBL1 and RUVBL2 are also clinically relevant and targetable dependency genes in NB. Only 2 studies hinted towards a role of RUVBL in NB previously: *Zhan et al.* reported on the involvement of RUVBL2 in mediating SK-N-DZ NB cell death upon histone deacetylase inhibition^44^, while very recently, *RUVBL1* was found to be part of a multigene prognostic signature in NB^45^. Interestingly, our results suggest that the apoptosis induction upon RUVBL inhibition is mediated by p53 and non-p53 pathways. Indeed, while we only found a transcriptomic signal of p53-induced apoptosis in CLB-BAR cells and not SK-N-AS cells (*TP53* is deleted in these cells^46^), all cell types examined responded with a clear induction of apoptosis and decreased cell growth upon experimental RUVBL inhibition.

In conclusion, our study has identified *RUVBL1* and *RUVBL2* as clinically relevant and therapeutically targetable dependency genes with independent prognostic value in NB. Our results suggest critical interactions between *RUVBL1/2, MYC(N)* and *ATR* and highly converging downstream signaling pathways in NB. While this opens several options towards novel combinatorial treatment strategies, one of the key future challenges will be to identify the most efficacious and least toxic therapeutic strategies in preclinical and later clinical studies.

## DATA AVAILABILITY STATEMENT

Raw RNA-seq data (fastq files) are available in ArrayExpress (https://www.ebi.ac.uk/arrayexpress/, accession nr: E-MTAB-13137). Quantified data are interactively available in the CLEAN web application at https://ccgg.ugent.be/shiny/clean/siaw_2024/^32^. All CUT&RUN data are available through the GEO repository (accession nr: GSE246722). All other data required to evaluate the conclusions in the paper are provided in Supplementary information files. Source code used for RNA-Seq and public data analyses is available at GitHub https://github.com/CCGGlab/RUVBL.

## METHODS

### Cell culture and CB-6644 drugging

Eight different NB cell lines were used in this study. CLB-BAR (*MYCN*-amplified), CLB-GE (*MYCN*-amplified) and CLB-GA (*MYCN*-nonamplified) cells were obtained from The Centre Leon Berard, France under MTA. SK-N-AS (*MYCN*-nonamplified), SH-SY5Y (*MYCN*-nonamplified) and SK-N-BE(2) (*MYCN*-amplified) cells were purchased from ATCC. NB1 cells were purchased from JCRB (Japanese Cancer Research Resources Bank). SHEP-MYCN cell lines were kindly provided by Tanmoy Mondal (University of Gothenburg, Sweden). All cell lines were tested for mycoplasma. Cell lines were cultured in complete media, RPMI 1640 supplemented with 10% foetal bovine serum (FBS) and a mixture of 1% penicillin/streptomycin at 37L°C and 5% CO_2_.

For CB-6644 IC_50_ determination, cells were seeded in 96-well plates at densities between 3000 and 5000 cells/well, and then treated with increasing concentrations of CB-6644. Cell viability was assessed after 3 days using resazurin assay.

Cell confluency/proliferation was monitored live using the IncuCyte® S3 Live Cell Analysis system (Essen BioScience) for 5 days (after treatment). Rate of cell growth under all conditions were determined using the IncuCyte® S3 software.

### Foci formation assay

Cells (1.0 × 10^5^) were seeded in 6-well plates and cultured overnight prior to treatment with 250 nM CB-6644 for 14 days. Cells were washed in PBS and fixed with methanol, followed by staining with 0.2% crystal violet, and washed. Plates were then scanned using Toshiba Studio 2505AC.

### Immunoblotting analyses

To evaluate the effect of CB-6644 or *MYCN* induction on RUVBL1/2 and downstream protein expression, NB cell lines (CLB-BAR, CLB-GA, NB1, SK-N-AS and/or SHEP-MYCN) were treated with 250 nM CB-6644 for 24h, 48h and 72h. Protein lysates were collected by lysing cells in RIPA lysis buffer and protein concentration was measured by BCA assay. Samples were subjected to western blot analyses. Chemiluminescence detection was done using Odyssey Fc Imager (LI-COR).

### Cell cycle analysis

SK-N-AS and CLB-GA (0.65 – 1 million cells) were seeded in a T-25 flask. After overnight culturing, cells were treated for 48h with CB-6644 at IC50 concentrations (SK-N-AS: 250 nM; CLB-GA: 120 nM) or DMSO control. Cells were washed once with ice-cold PBS before collection in Phosphate-buffered saline (PBS). During each wash step, cells were centrifuged for 5 minutes at 1200 rpm and supernatant was removed. Cells were washed once in ice-cold PBS before fixation in 70% ethanol for at least 1 hour on ice, then washed once with ice-cold PBS and incubated in PBS with ribonuclease A (RNase A, 250 µg/mL) for 1 hour at 37°C. Propidium iodide (40 µg/mL) was added, and analysis was performed on a BD LSR II flow cytometer using BD FACSDIVATM Software. The flow cytometry results were analysed using FlowJoTM v10.8 Software (BD Life Sciences).

### Apoptosis Assay

Cells (7000 per well) were seeded in 96-well plate and cultured overnight. They were then treated with 250 nM CB-6644 for either 48 hrs or 72 hrs in three technical replicates. DMSO was used as negative control. Apoptosis was evaluated by measuring caspase 3 and 7 activities in the treated cells, using the Caspase-Glo® 3/7 Assay kit (Promega), following the manufacture’s protocol. Briefly, Caspase-Glo reagents were added to the cells in 96-well plate, incubated for 45 minutes with gentle shaking. Luminescence was recorded with the GloMax® system (Promega).

### siRNA-mediated knockdown of RUVBL1 and RUVBL2

NB cells were transfected with Silencer select siRNAs (Life Technologies) against *RUVBL1* [ID # S16369; CGAGUGAUGAUAAUCCGGAtt (siRUVBL1 #1) and S16370; GAAGUUUACUCAACUGAGAtt (siRUVBL1 #2)] and *RUVBL2* [ID # S21307; GGAGAUCCGUGAUGUAACAtt (siRUVBL2 #1) and S21309; GAAACGCAAGGGUACAGAAtt (siRUVBL2 #2)] using lipofectamine RNAiMAX transfection reagent (# 13778150, ThermoFisher Scientific). Scrambled siRNA (# S103650325, Qiagen) was used as negative control. Cells were harvested after overnight transfection and seeded (2.5 × 10^3^ to 4 × 10^3^) in 96-well plates. The remaining cells were seeded in 6-well plates and cultured for 5 days and lysed in RIPA buffer for western blot analysis. The 96-well plates were monitored for cell proliferation using the IncuCyte® S3.

### Organoid growth assay

Human high-risk NB PDX (NEC005) tumoroid, with *MYCN*-amplification, Chr 2p and 17q gains, and Chr 1p and 3p deletions, were dissociated into single cells using StemPro Accutase (# A11105-01; Gibco. Cells were then seeded at 3 different densities, 3000, 5000 and 7000 cells per well of 384 well plate. After overnight culture, cells were treated with either 250 nM or 500 nM CB-6644 in triplicates. DMSO was used as negative control. Organoid growth was monitored live with IncuCyte® S3.

### MYCN CUT&RUN analysis

CUT&RUN coupled with high-throughput DNA sequencing was performed on isolated nuclei using Cutana pA/G-MNase (Epicypher, 15-1016) according to the manufacturer’s manual. Briefly, nuclei were isolated from 0.5 × 10^6^ cells/sample in 100 µl nuclear extraction buffer per sample and incubated with activated Concanavalin A beads for 10 min at 4°C while rotating. Nuclei were resuspended in 50 µl antibody buffer containing a 1:100 dilution of N-MYC antibody (# sc-53993, Santa Cruz, RRID:AB_831602) or control IgG (#3900; CST, RRID:AB_1550038), and kept in an elevated angle on a nutator at 4°C overnight. Next, targeted digestion and release was performed with 2.5 µL Cutana pA/GMNase (15-1116) and 100 mM CaCl2 for 2 hours at 4°C on the nutator. After chromatin release by incubation on 37°C for 10 minutes, DNA was purified using the CUT&DNA purification kit (14-0050) and eluted in 12 µl of elution buffer. Sequencing libraries were prepared with the NEBNext Ultra II kit (Illumina, E7645), followed by paired-end sequencing on a Nextseq2000 using the NextSeq 2000 P2 Reagents 100 Cycles v3 (Illumina, 20046811). Prior to mapping to the human reference genome (GRCh37/hg19) with bowtie2 (v.2.3.1), quality of the raw sequencing data of CUT&RUN was evaluated using FastQC and adapter trimming was done using TrimGalore (v0.6.5). Quality of aligned reads were filtered using min MAPQ 30 and reads with known low sequencing confidence were removed using Encode Blacklist regions. For sample with a percentage duplicated reads higher than 10%, deduplication was performed using MarkDuplicates (Picard, v4.0.11). Peak calling was performed using MACS2 (v2.1.0) taking a q value of 0.05 as threshold and default parameters. Homer (v4.10.3) was used to perform motif enrichment analysis, with 200 bp around the peak summit as input. The R package *igvR* (v1.19.3) was used for visualization of the data upon RPKM normalization.

### Immunohistochemical staining of human NB samples

Human NB (n=3), and pancreatic tissues (n=2, control) were fixed in 10% neutral buffered formaldehyde and embedded in paraffin. Stainings for RUVBL1 and RUVBL2 were performed on 3-µm-thick sections via an automatic immunostainer (BenchMark Ultra, Ventana Medical Systems). Two anti-RUVBL1 rabbit polyclonal antibodies (HPA019948, Sigma-Aldrich: 1/100, incubation 32 min and HPA019947, Sigma-Aldrich: 1/500, incubation 32 min) and 2 anti-RUVBL2 rabbit polyclonal antibodies were used (HPA067966, Sigma-Aldrich: 1/500, incubation 32 min and PA5-29871, ThermoFisher Scientific: 1/50, incubation 32 min). Visualization was achieved with the Ultraview Universal DAB Detection Kit (Ventana Medical Systems). Heat-induced epitope retrieval was performed with Cell Conditioning 1 (Ventana Medical Systems), 64 min at 95°C, except for anti-RUVBL1 antibody HPA019948 for which an incubation time of 36 min was used.

### Data download and processing

Human NB RNA-seq (batch corrected log normalized counts) and related clinical data were obtained from the study of *Cangelosi et al.*^47^ (available at https://www.ncbi.nlm.nih.gov/pmc/articles/PMC7563184/bin/cancers-12-02343-s001.zip).

RNA-Seq differential gene expression data from *Th-MYCN*-driven NB mice models treated with the ATR inhibitors elimusertib or ceralasertib (both 25 mg/kg) were obtained from our previous studies^19,21^ (available at https://static-content.springer.com/esm/art%3A10.1038%2Fs41467-021-27057-2/MediaObjects/41467_2021_27057_MOESM3_ESM.xlsx and https://www.pnas.org/doi/suppl/10.1073/pnas.2315242121/suppl_file/pnas.2315242121.sd02.xlsx). RNA-Seq data from MYCN-induced SHEP cells and elimusertib-treated CLB-BAR cells were retrieved using CLEAN (https://ccgg.ugent.be/shiny/clean/)^32^.

Gene expression profiling microarray data from *MYCN*-driven NB tumor development in the *TH-MYCN* mouse model were obtained from ArrayExpress (accession nr. E-MTAB-3247)^48^. Background correction and quantile normalization of these data were performed using the *Limma* v.3.50.3 *R* package^49^. Gene expression of *Ruvbl1*, and *Ruvbl2* was evaluated at 1, 2, and 6 weeks. Genes differentially expressed between *Th-MYCN*^+/+^ and wild-type mice at week 6 were determined using *Limma* moderated t-statistics to determine significance in gene expression changes.

Gene effect scores (dependency scores) of 34 NB cell lines, derived from CRISPR knockout screens and published by Broad’s Achilles and Sanger’s SCORE projects, were downloaded from DepMap^30^ (https://depmap.org/portal/download/all/). Negative gene effect scores imply cell growth inhibition and/or death following gene knockout. Non-essential genes have a normalized median score of 0 and predefined common essential genes have a median score of -1. Essential genes (n = 1910) were derived from DepMap.

### Gene set enrichment analysis (GSEA)

Preranked GSEA was performed using the R *fgsea* package (*fgseaMultilevel* function, default parameters) with ranking based on the DEseq2 statistic. Mouse (GTRD transcription factor targets) and human (Hallmark, Reactome) gene sets were downloaded from the Molecular Signatures Database v2023.1.

### RNA-Sequencing and quantitative PCR

CLB-BAR and SK-N-AS NB cell lines were treated with 250 nM CB-6644 for 24, 48 and 72 hrs. DMSO was used as negative control. RNA was extracted from the cell pellets using the ReliaPrep™ RNA Miniprep Systems (Promega) and the manufacturers protocol was followed. RNA samples were either used for RNA-seq or quantitative PCR.

RNA sequencing was performed by Biomarker Technologies (BMK, Germany). RNA-Seq paired-end reads (read length 150 base pairs) were aligned to the GRCh38 reference genome using *HISAT2*^50^. The average alignment efficiency for all samples was 93.4%. Genes were annotated using GENCODE 29 and quantified using *HTSeq*^51^. Further analysis was performed using only coding genes. Differential gene expression was determined using *DESeq2*^52^. Only expressed genes, defined as genes with a basemean value higher than 10, were considered for further analysis. Genes were considered differentially expressed if their absolute log2 fold change values were above 1 at FDR-adjusted p values (*Padj*) below 0.05.

cDNA was synthesized using the iScript cDNA synthesis kit (Biorad) and quantitative PCR was performed on StepOnePlus Real-Time PCR Systems using Power SYBR® Green master mix and following primers: MYCN: 5’- CTGAGCGATTCAGATGATGAAG-3’ and 5’-CCACAGTGACCACGTCGATT-3’; MYC: 5’-CAGCTGCTTAGACGCTGGAT-3’ and 5’-AGCTAACGTTGAGGGGCATC- 3’; ACTB: 5’-ATGACCCAGATCATGTTTGAGAC-3’ and 5’- CCAGAGGCGTACAGGGATAG-3’.

### Survival analysis

Overall survival (OS) analyses were performed using the R packages *Survival v.3.4-0* and *Survminer v.0.4.9*. OS curves were plotted by the Kaplan-Meier method. Tumor samples were stratified (high vs low gene expression) based on median gene (*RUVBL1* or *RUVBL2*) expression. Log-rank tests were performed to assess statistical significance.

The multivariate Cox proportional hazards regression model was used to evaluate the prognostic values of genes. Tumor stage (stage 4 vs stages 1,2,3), age at diagnosis (>18 months vs <18 months) and *MYCN* amplification were used as covariates in the multivariate analysis.

### Data analysis and statistics

Statistical analyses were performed with R statistical package. Statistical tests are indicated in the respective sections and figure captions. Multiple testing corrections were performed using the Benjamini-Hochberg method^53^.

## SUPPLEMENTARY FIGURE LEGENDS

**Supplementary figure 1.**
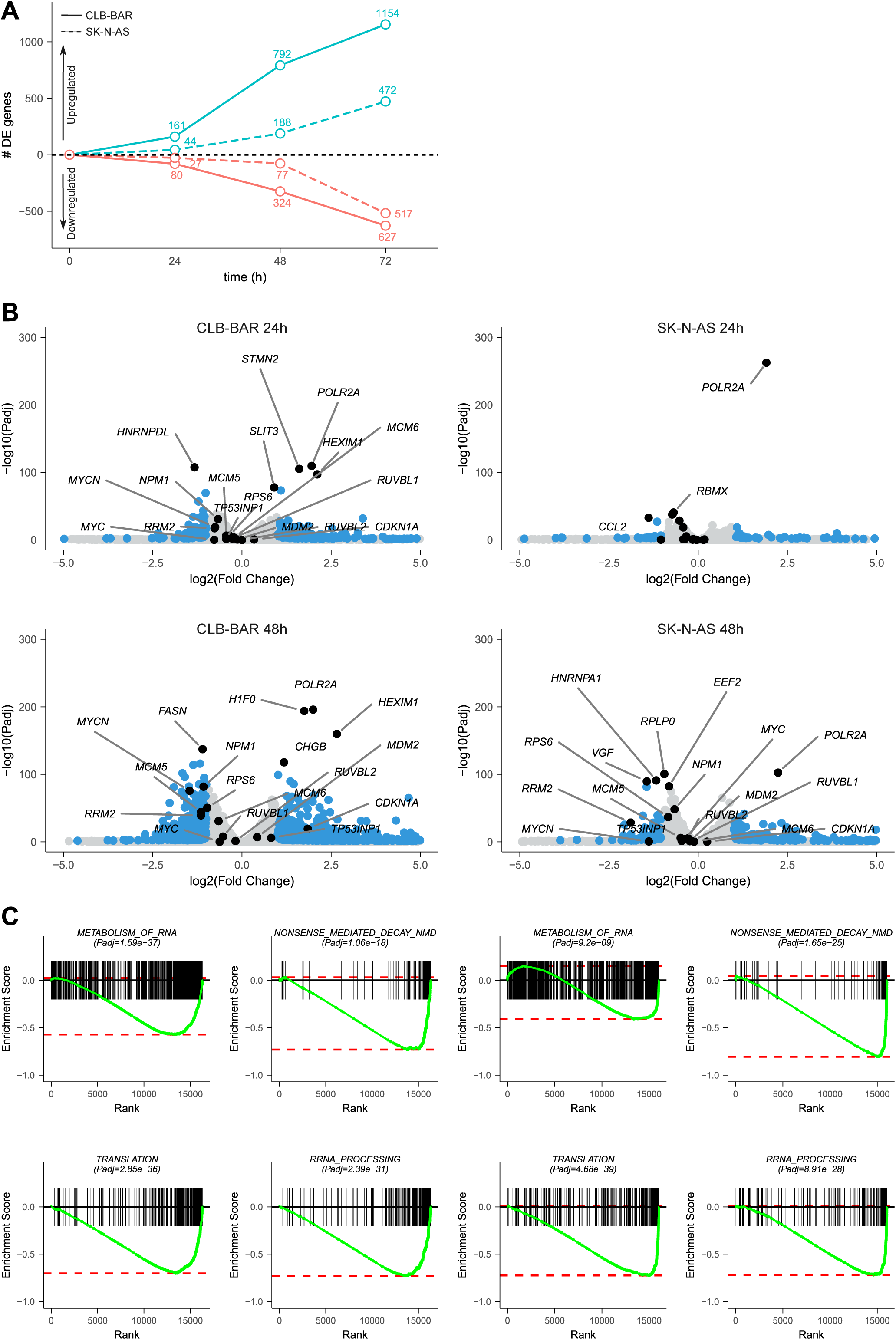
Transcriptomic response to CB-6644 treatment of NB cells. CLB-BAR and SK-N-AS NB cells were treated for 24-72h with CB-6644 250 nM and differential gene expression (DGE) was determined. **A.** Number of up- and downregulated genes (threshold log2FoldChange of ±1 at 1% FDR) for both cell lines and 3 time points as indicated. **B.** Volcano plots showing DGE results for both cell lines after 24h and 48h of treatment as indicated. Differentially expressed genes indicated in blue with top up/downregulated and genes discussed in main text labelled. **C.** GSEA running score plots for 4 Reactome gene sets in both cell lines as indicated. See Suppl. Table 1 for detailed DGE and GSEA results.

**Supplementary figure 2.**
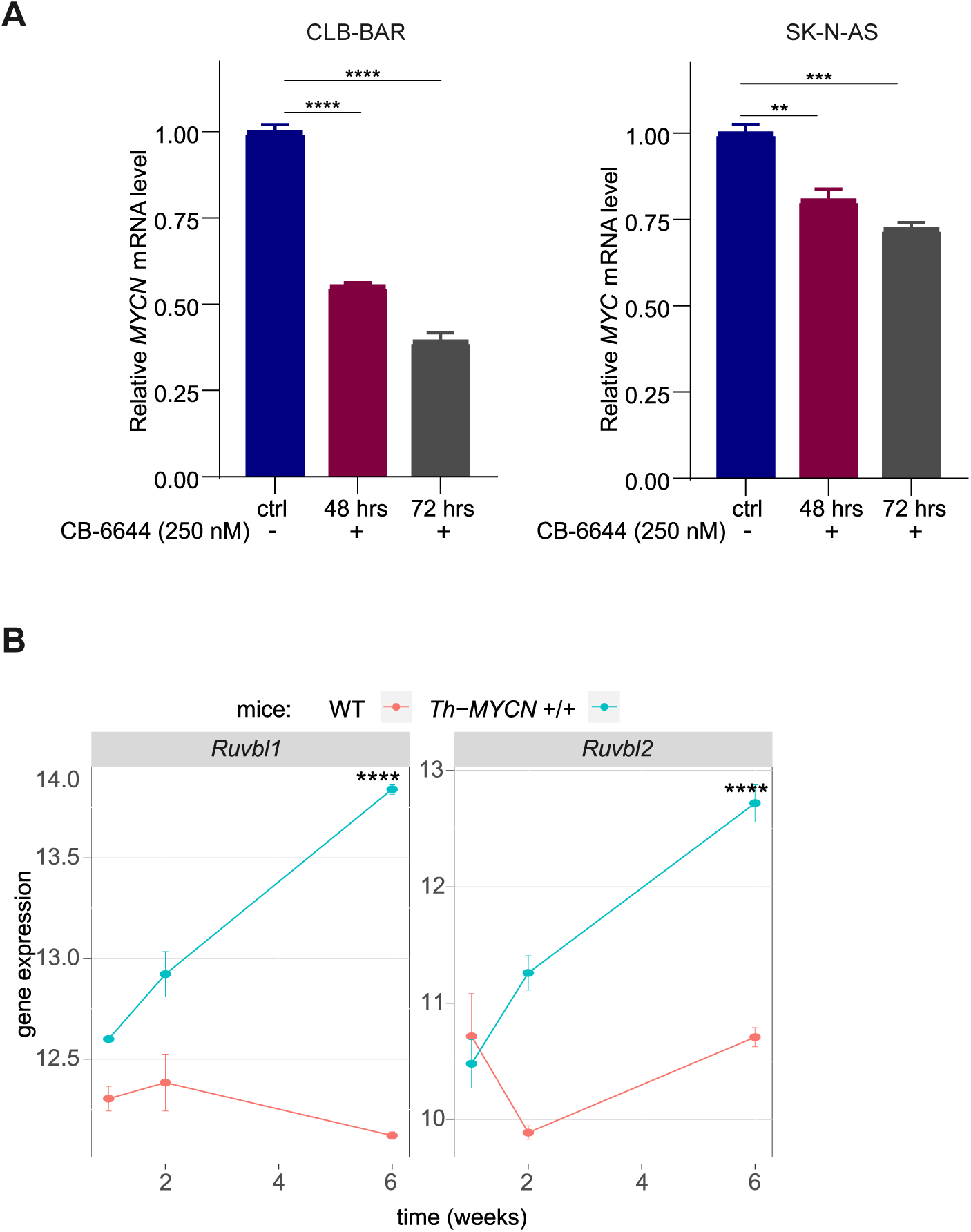
RUVBL1/2 and MYCN correlation in cell lines and MYCN-driven mice models. **A.** Quantitative PCR was performed after treatment of CLB-BAR and SK-N-AS cells with CB-6644 (250 nM) for 48h and 72h. Bar plots indicating *MYCN* (CLB-BAR) and *MYC* (SK-N-AS) mRNA levels. Results are means +/-SEM of three independent biological replicates. ** *P* value < 0.01, *** *P* value < 0.001, **** *P* value < 0.0001. Unpaired, two-sided Student’s t-test **B.** Time course of *Ruvbl1* and *Ruvbl2* gene expression during control (WT, wild-type) and Th-MYCN-driven tumorigenesis in mice. Data derived from *De Wyn et al*.^48^ ****, Padj < 0.0001, moderated t-statistic.

**Supplementary figure 3.**
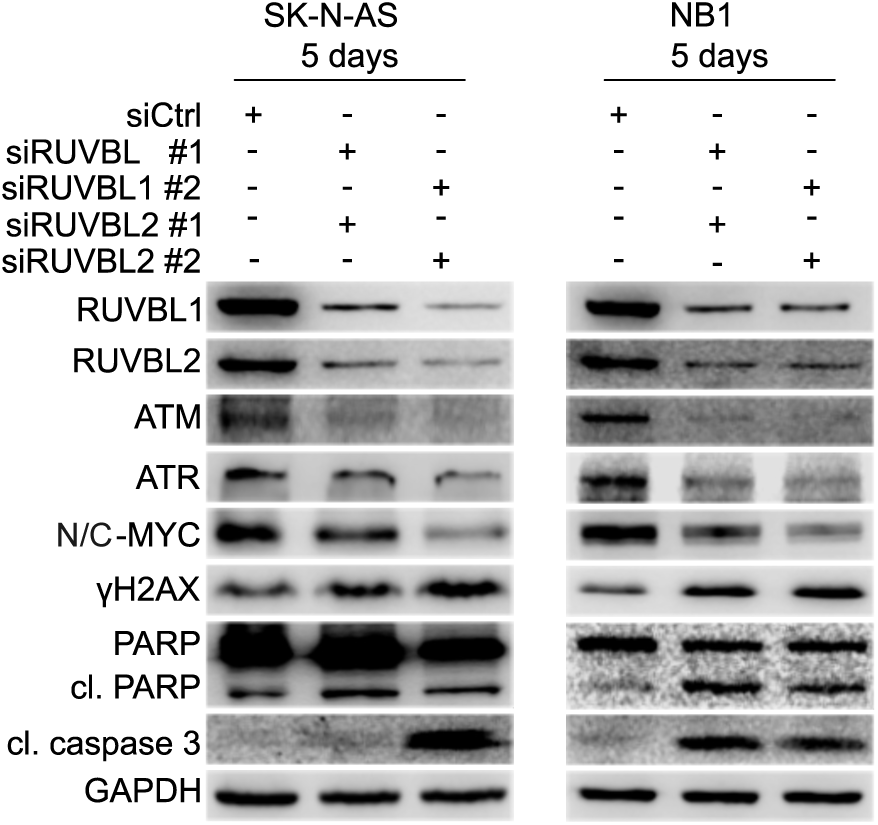
Effect of siRNA-mediated knockdown of *RUVBL1* and *RUVBL2* in NB cell lines. Western blot showing the effect of siRNA-mediated *RUVBL1* and/or *RUVBL2* knockdown on downstream signaling proteins, 5 days after transfection of two independent NB cell lines as indicated. Blots are representative of two independent experiments. Cl, cleaved.

**Supplementary figure 4.**
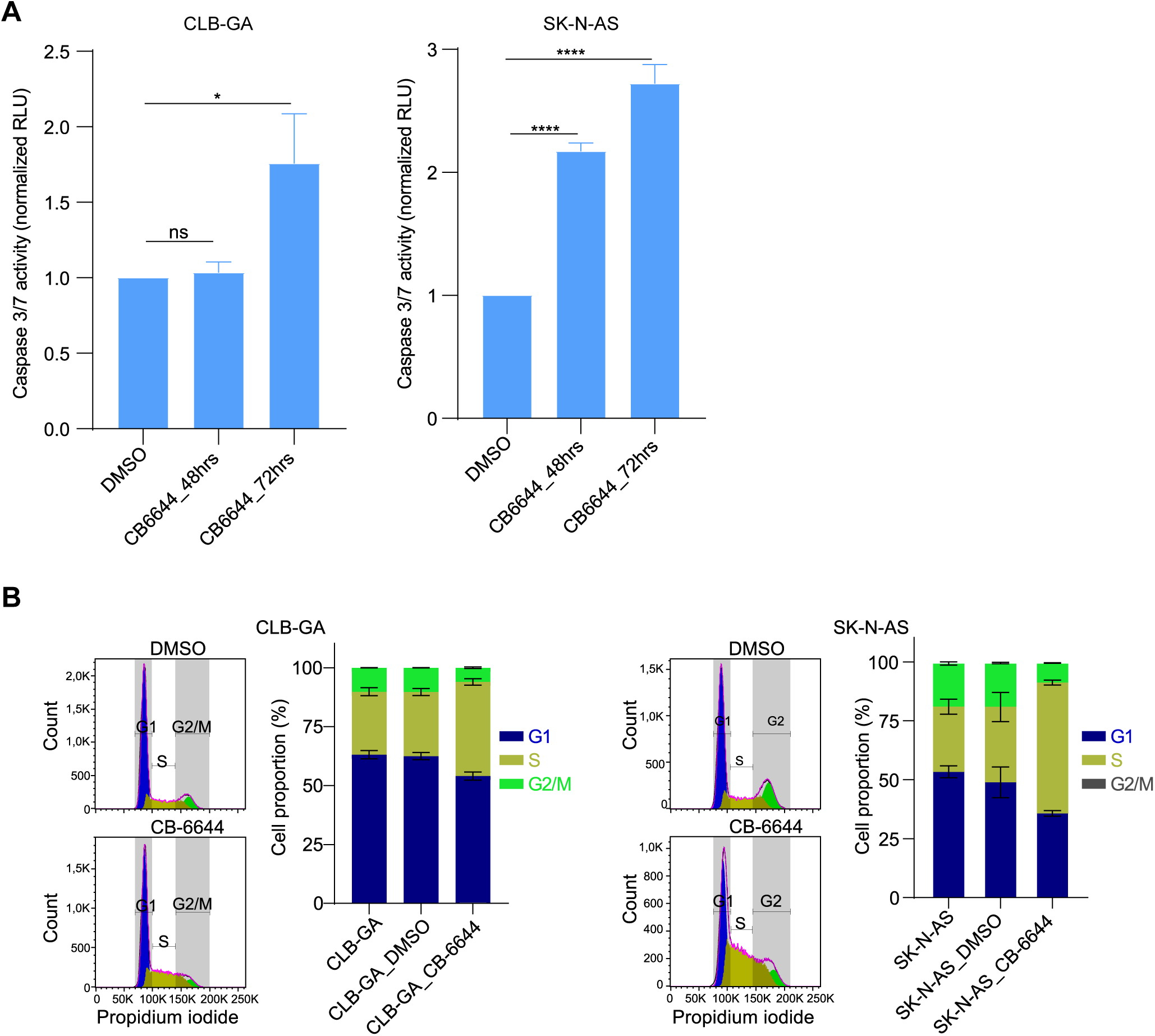
Experimental validation of apoptosis induction and cell cycle alterations upon NB cell treatment with CB-6644. **A.** CLB-GA and SK-N-AS NB cells were treated with CB-6644 (250 nM) and caspase 3/7 activity was measured after 48h and 72h as indicated in bar plots. Caspase activity was quantified using relative light unit (RLU) and normalized to DMSO (control) groups. Results are means +/-SEM of three independent biological replicates. ns: not significant, * *P* value < 0.05, **** *P* value < 0.0001, unpaired two-sided t-test. **B.** Flow cytometry-based cell cycle analyses of propidium iodide-stained NB cell lines treated for 48h with DMSO or CB-6644 at IC50 concentrations (CLB-GA: 120 nM; SK-N-AS: 250 nM). Stacked bar plots show proportions in each cell cycle phase, at different conditions, for two experimental repeats

## AUTHOR CONTRIBUTIONS

This study was conceptualized and designed by JTS and JVdE. qPCR analyses and cell viability assays of mouse NB cells were performed by W-YL and MB. Cell cycle, organoid and CUT&Run experiments were conducted by EH, ES and SB and supervised by FS and KD. Immunohistochemical staining was performed by IK and JVD. JTS conducted all other wet-lab experimental analyses. JTS, AC and JVdE performed the transcriptomics data analyses. JTS, RHP and JVdE drafted the manuscript, with subsequent review and editing from all authors.

## ACKNOWLEDGEMENT

This project was funded by grants from the Swedish Childhood Cancer Foundation (TJ2021-0068 - JTS; PR2022-0029 - RHP; PR2021-27 - BH), the Swedish Research Council (R.H.P.:2019-03914; B.H.:2021-01192), the Assar Gabrielsson’s foundation (FB22-24 - JTS), Cancer Research Institute, Ghent (YIPOC-2023 - JTS), the Ghent University Special Research Fund (BOF.STG.2019.0073.01 - JVdE; BOF.GOA.2022.0003.03 - FS), Kom op tegen Kanker (Stand up to Cancer), the Flemish cancer society (STI.VLK.2022.0013.01 - AC), the Swedish Cancer Society (CAN21/1459 - RHP; CAN21/1525 - BH), Villa Joep grant (FS) and the Research Foundation Flanders (FWO; FWO.OPR.2023.0063.01 - FS, JVdE, RHP and V424522N to JVdE).

